# Variation analysis of SARS-CoV-2 complete sequences from Iran

**DOI:** 10.1101/2021.01.23.427885

**Authors:** Jale Moradi, Mohsen Moghoofei, Amir Houshang Alvandi, Ramin Abiri

## Abstract

The SARS-CoV-2 is a new emerging coronavirus initially reported in China at the late December 2019 and rapidly spread to the whole of the world. To date, 1261903 total case and 55830 deaths are reported from Iran as 8 January. In this study, we investigated all the complete sequences of SARS-CoV-2 that publicly reported from Iran. Twenty-four sequences between March to September 2020 were analyzed to identify genome variations and phylogenetic relationships. Furthermore, we assessed the amino acid changes related to the spike glycoprotein as an important viral factor associated with the entry to the host cells and as a vaccine target. Most of the variations are occurred in the ORF1ab, S, N, intergenic and ORF7 regions. The analysis of spike protein mutations demonstrated that D614G mutation could be detected from the May and beyond. Phylogenetic analysis showed that most of the circulated viruses in Iran are belong to the B.4 lineage. Although, we found a limited number of variants associated to the B.1 lineage carrying D614G mutation. Furthermore, we detected a variant characterize as the B.1.36 lineage with sixteen mutations in the spike protein region. This study showed the frequency of the viral populations in Iran as September, therefore, there is an emergent need to genomic surveillance to track viral lineage shift in the country beyond the September. These data would help to predict future situation and apply better strategy to control of the pandemic.

## Introduction

Severe acute respiratory syndrome coronavirus 2 (SARS-CoV-2) is a new emerging single-stranded RNA virus which initially was reported in Wuhan, Hubei Province, China at the late December 2019 (1). The disease was rapidly spread to other countries. According to the WHO, there were 85,929,428 confirmed cases and 1,876,100 confirmed deaths reported until 8 January, 2021.

The SARS-CoV-2 consists of seven canonical open reading frames (ORFs) including ORF1ab, ORF3a, ORF6, ORF7a, ORF7b, ORF8 and ORF10. These ORFs encode four structural proteins including spike (S) glycoprotein, envelope (E), membrane (M), nucleocapsid (N) proteins, nine accessory proteins and 16 nonstructural proteins (nsp1-16) (2). The fast spread of the virus throughout the world with various hosts and environments may lead to rise of different virus populations due to facing different selective pressures. Thus, many studies are performed to identify virus variations in different geographical regions (3,4). Some viral proteins are targeted by the immune system that variations in these parts of the genome would affect the virulence and transmissible potential (5). Furthermore, mutations can interrupt the vaccines efficacy and validity of diagnostic tests due to the changes in the targeted proteins and probes (6,7).

Now, there are 290,997 complete genome sequences of the SARS-CoV-2 in GISAID and the number of the submitting sequences are increasing. There is no classifying system for SARS-CoV-2 variants in International Committee on Taxonomy of Viruses (ICTV). Although, many scientists are trying to characterize the genetic diversity of the variants. In the early of the pandemic, two clades named S and L were introduced, and then L evolved as another clade. Until recently, there are six clades circulating in the world based on the sequences submitted in the GISAID (4). In another nomenclature, eighty-one lineages were identified for SARS-CoV-2 phylogeny belong to the A and B lineages (8).

There are large number of SARS-CoV-2 genome sequences that are deposited in the GISAID public database, and it is possible to characterize the evolution pattern of the virus geographically by phylogenetic analysis (9).

## Methods

In total, 25 SARS-CoV-2 complete sequences related to Iran were released by November 25, 2020. One of the retrieved sequences was removed due to a high N content (more than 5%). All of 24 sequences were aligned to SARS-CoV-2 reference genome (NC_045512.2) using MAFFT (v7.455) (10). We applied SNP-sites to extract the genome variations from the multiple sequence alignment (MSA) file (11). Then, the association of each variation to SARS-CoV-2 ORFs was surveyed (2). Furthermore, we analyzed the amino acid changes related to the spike protein. For phylogenetic analysis, the MSA file was visualized for checking the quality and trimming using the UGENE software (12). The trimmed file was applied to construct phylogenetic tree with maximum likelihood method using RaxML-NG v.0.9.0 (applying 1000 bootstrap) (13). The constructed tree was visualized using FigTree v1.4.4.

## Results

### Detection of genome variation sites

In total, 24 complete sequences were analyzed, which 11 of the sequences were reported from Tehran (capital of Iran), 1 from Semnan (east of Iran), 1 from Qom (center of Iran), 1 from Zahedan (south of Iran) and 9 from unknown sources (Supplementary table 1). In total, 275 variation sites were detected that 191 variation sites were found in ORF1ab and 32, 13 and 10 were related to S, N and intergenic regions, respectively (Supplementary Table 2). The other genome segments had less than 10 variations sites.

### Frequency of variations in the samples

Of the 419 variations that were observed in the 24 sequences, most of the variants were related to the 21 variation sites (Table 1). Total of 278 variations were related to the ORF1ab and 44, 33, 31 and 11 variations were detected in S, N, intergenic and ORF7 regions, respectively (Fig. 1). Our analysis showed that there is a uniform frequency of variations among the samples. Variations had occurred in the 5 to 15 points in most of the sequences, although there were three sequences with 203, 23 and 25 variation sites (Fig. 2). The collection date of the sample was from March to September of 2020, 14 of the sequences were collected in March, 1 were September and 3 sequences were collected for each three months as April, May and June. The mutations occurred in the steady rate except the May that two samples with high mutation quantity were seen (203 and 23 mutations) (Fig. 3).

**Figure 1:**
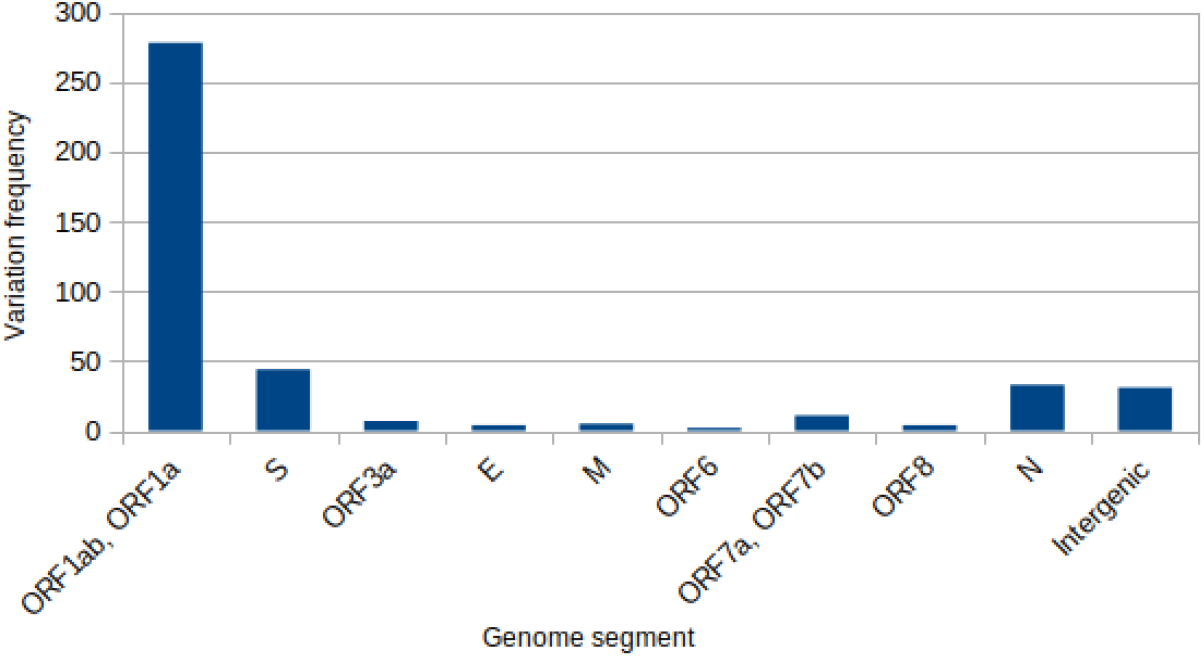
Frequency of the nucleotide variations in SARS-CoV-2 genomes

**Figure 2:**
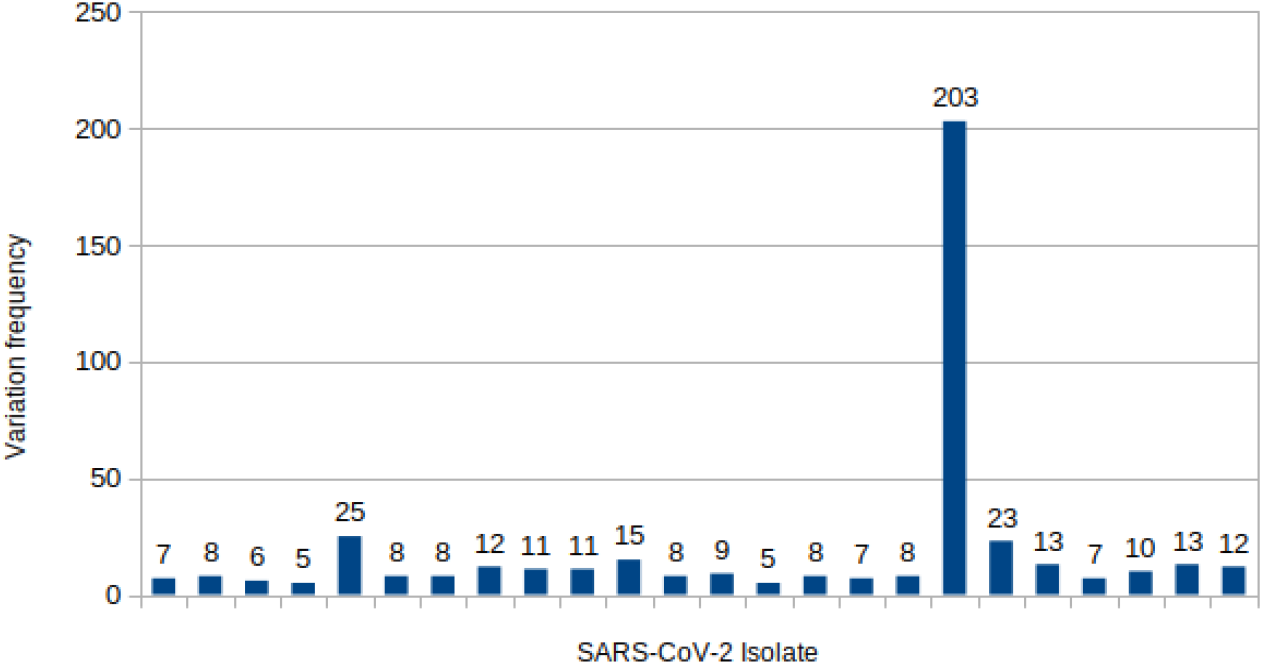
Frequency of the nucleotide variations in SARS-CoV-2 isolates

**Figure 3:**
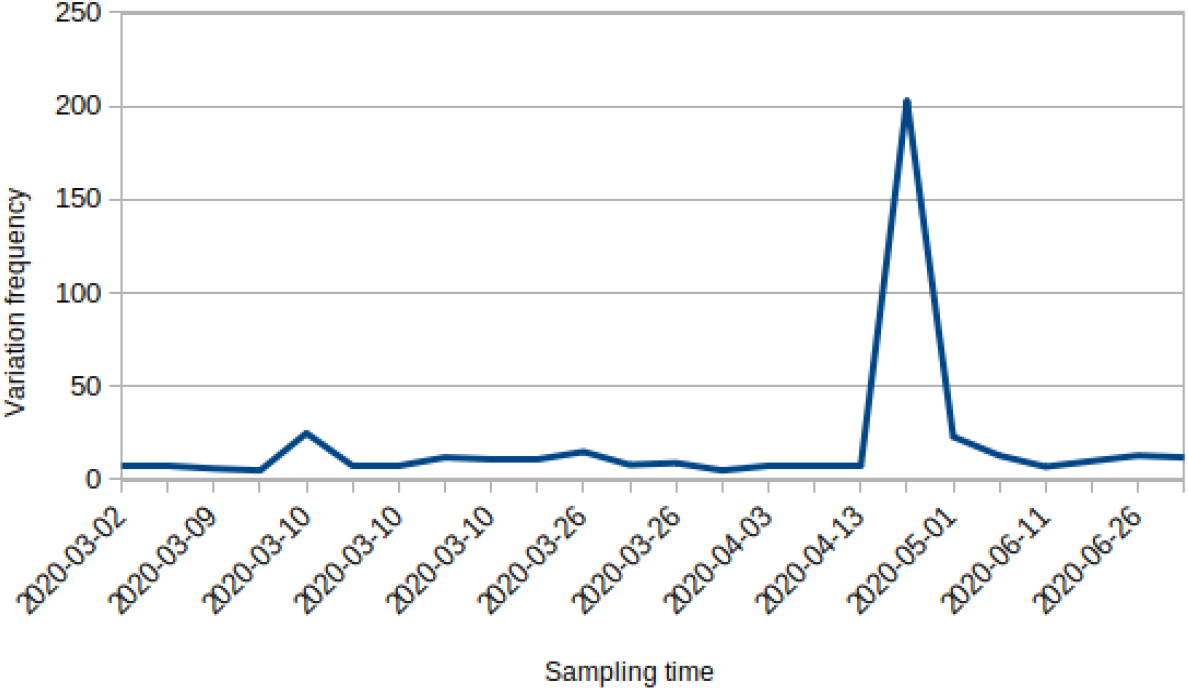
Frequency of the nucleotide variations in SARS-CoV-2 sampling time

**Table 1:** Prevalent nucleotide variations in SARS-CoV-2 sequences

| Position | Variation Frequency | Reference | Alternative | ORF |
| --- | --- | --- | --- | --- |
| 241 | 5 | C | T | ORF1ab, ORF1a |
| 517 | 3 | TATG | T | ORF1ab, ORF1a |
| 884 | 8 | C | T | ORF1ab, ORF1a |
| 1397 | 19 | G | A | ORF1ab, ORF1a |
| 3037 | 5 | C | T | ORF1ab, ORF1a |
| 8653 | 8 | G | T | ORF1ab, ORF1a |
| 11083 | 15 | G | T | ORF1ab, ORF1a |
| 14408 | 4 | C | T | ORF1ab |
| 15842 | 3 | C | T | ORF1ab |
| 17288 | 3 | C | T | ORF1ab |
| 18555 | 3 | C | T | ORF1ab |
| 20887 | 4 | G | A | ORF1ab |
| 21627 | 4 | C | T | S |
| 22735 | 3 | C | T | S |
| 23277 | 3 | C | T | S |
| 23403 | 4 | A | G | S |
| 27788 | 3 | G | T | ORF7b |
| 28688 | 17 | T | C | N |
| 28830 | 3 | C | T | N |
| 28854 | 3 | C | T | N |
| 29742 | 18 | G | T | Intergenic |

### Frequency of Spike glycoprotein mutations

Seventeen sequences had amino acid changes in the spike glycoprotein (Table 2). These sequences were sampled throughout the pandemic, although, we analyzed D614G mutation with or without other mutations A475V, L452R, V483A, and F490L that are related to the increasing virus infectivity and transmissibility potentials. We found that the sequences that were belonged to May and beyond had D614G mutation without any of the other important co-existent mutations. Furthermore, the sequence with 203 variation sites that described earlier, had 16 types of the spike mutations including D614G.

**Table 2:**
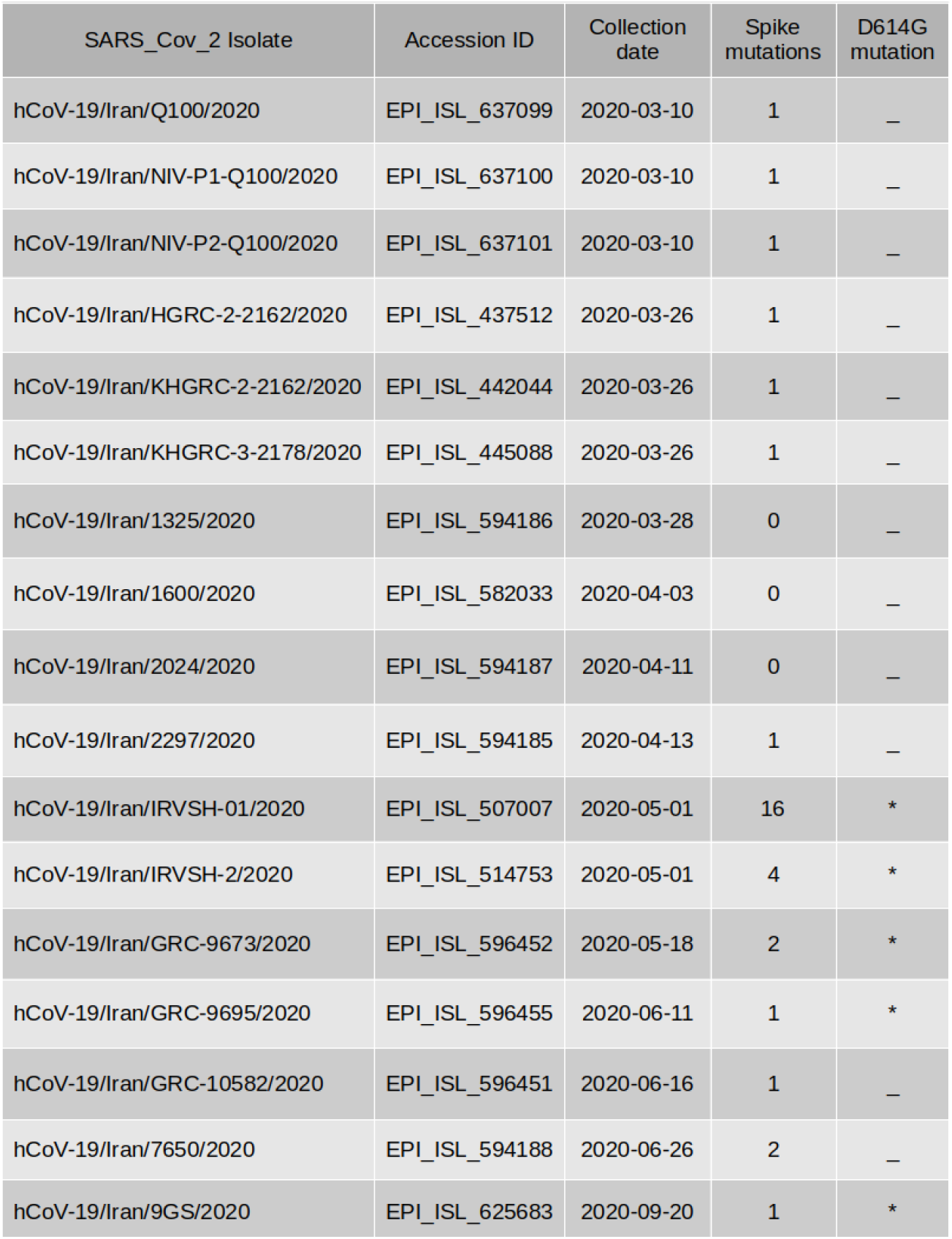
Frequency of Spike protein mutations in SARS-CoV-2 sequences

### Phylogenetic analysis

GISAID has introduced six phylogenetic groups of SARS-CoV-2 genomes represented by S and L groups at the first of the pandemic, followed by V and G groups which were evolved from group L. Groups G, GH and GR are most recent derivative ones. In another nomenclature system, SARS-CoV-2 is classified in 81 viral lineages named groups A and B (8). Totally, six lineages are grouped in lineage A named A.1 to A.6, among which lineage A.1 consist of two sub-lineages. Also, Lineage B includes 16 lineages that B.1 lineage is the most prevalent worldwide and consists more than seventy sub-lineages. In the current study, we analyzed the frequency of the detected lineages (Fig. 4). Although, the B.4 was the predominant lineage that 19 sequences are classified in this lineage type (Blue items, Fig. 5) while just four sequences are categorized in lineage B.1 (purple ones, Fig. 5). As It is demonstrated, four sequences that are belongs to B.1 sub-lineages are related to the clade G, and three sequences in lineage B.4 are related to the clade L, and the final sequences in B.4 lineage is not correlated to a GISAID clade. Furthermore, our analysis showed that all D614G mutations related to spike protein are related to the B.1 lineage sequences.

**Figure 4:**
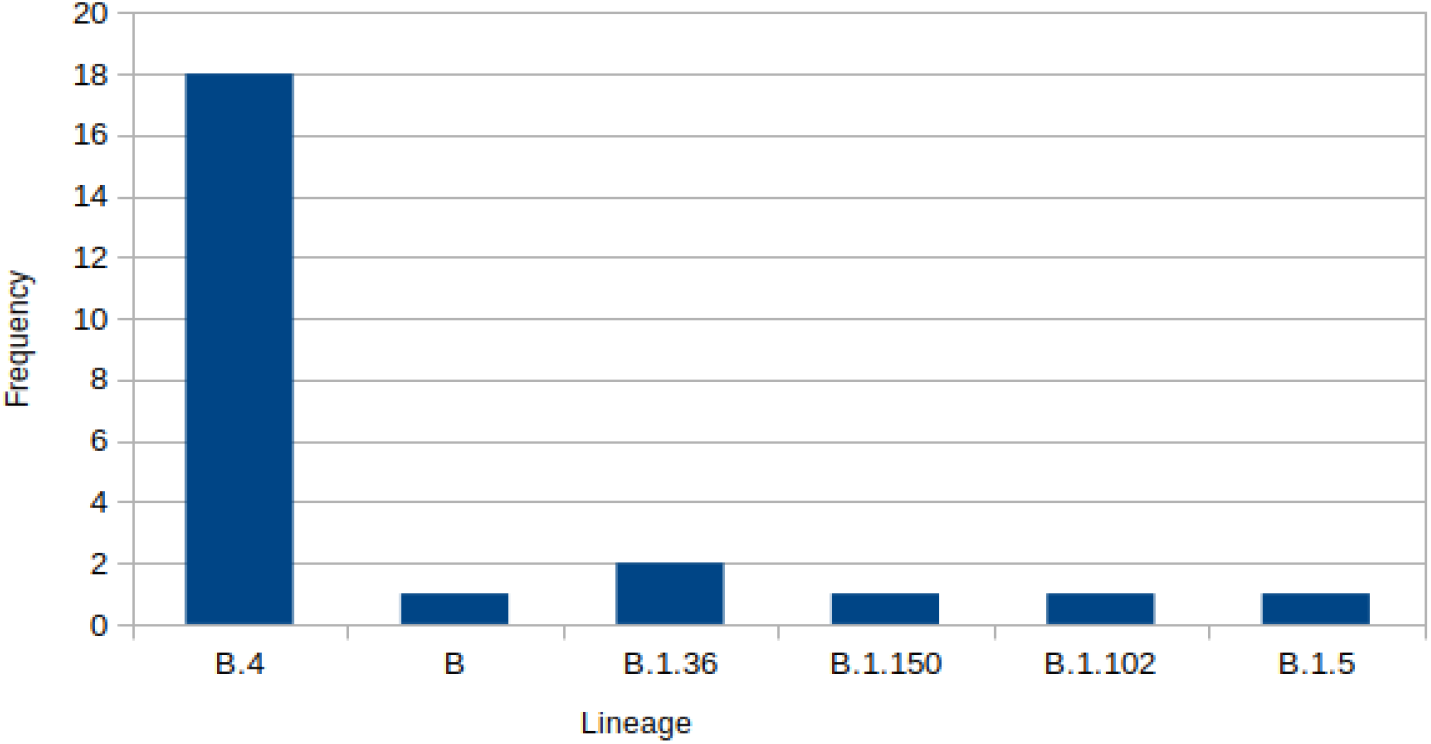
Frequency of SARS-CoV-2 lineages

**Figure 5:**
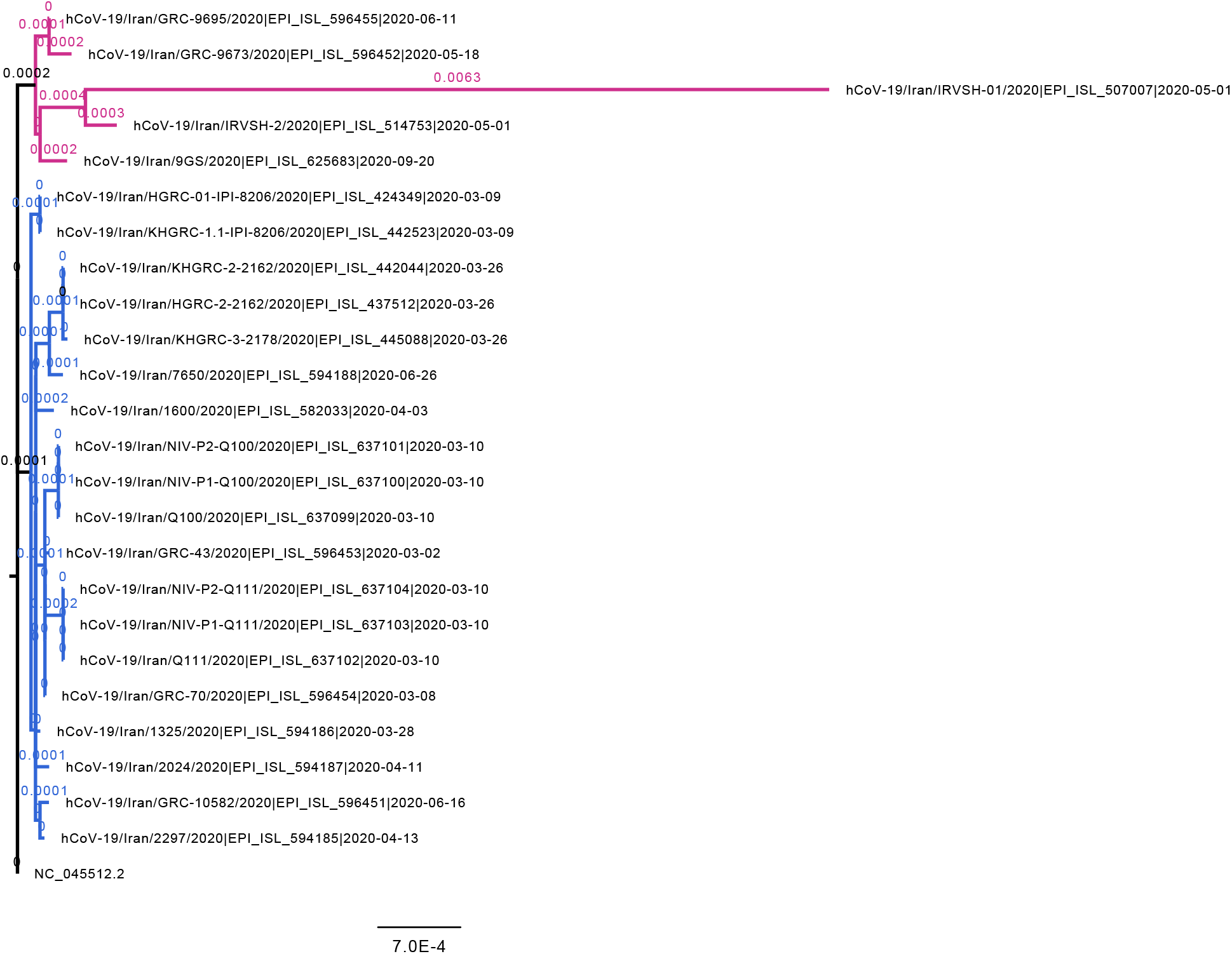
Phylogenetic analysis of SARS-CoV-2 sequences

## Discussion

The genome of SARS-CoV-2 includes seven major ORFs and 23 unannotated ORFs (2,14). ORF1ab consists of two overlapping ORFs (ORF1a and ORF1b) that occupied two-third of the viral genome and encodes a poly-protein which is cleaved to 16 nonstructural proteins. According to our results and many other reports, most of the variation sites are raised in ORF1ab. Other major ORFs encode four canonical structural proteins including spike (S), membrane (M), envelope (E) and nucleoside (N). Since the amino acid sequences must be conserved for an ideal immunogen protein (in this case, S protein), there is an urgent need to characterize the rate of the mutations in this part of the genome in all geographic regions (15,16). Our results demonstrated that approximately 10% of the variation sites belong to S protein, although, these result in the 34 amino acid changes. Nucleocapsid and intergenic regions are other mutation vulnerable sequences. Spike and N proteins have higher mutation rate after the ORF1ab in the other studies that is compatible with our results (4). However, mutations in the intergenic sites are much higher in our experience comparing the same studies.

The number of the variations were consistent among the isolate, although there was a sample with high mutation rate. This strain was isolated in May 2020 and showed 203 nucleotide variations. This variant is categorized to the lineage B.1.36 and previous studies showed that this lineage could contain various spike mutations which results in less reactogenicity to the antibodies generated by vaccine candidates (17). We found that this isolate has 16 types of the mutation in the spike glycoprotein including D614G. The Spike mutations were seen in isolates with different sampling time, although D614G mutations only existed in the specimens which were collected from May and after. In a systematic review, 80 variants and 26 glycosylation mutants of the spike protein were surveyed in the terms of infectivity and reactivity, results showed that most of the variants were susceptible to the neutralizing antibodies, although, D614G in coexistence with some variants including A475V, L452R, V483A, and F490L were unidentifiable by neutralizing antibodies and were more infectious (18–20). Spike protein’s D614G mutation was detected in March 2020 in 26% of the studied sequences, but its frequency increased to 74% by June 2020 (21). As mentioned earlier, we only detected D614G mutation without other infectivity related mutations. Totally, our analysis in the term of D614G mutation screening indicated that this mutation showed up in the Iran in May when the frequency of this variant had been increased globally. Plante et al. demonstrated that this mutant virus has more potential of viral replication in the human lung tissue. Also their report showed that D614G ability to S1/S2 cleavage substitution and shedding of spike protein had changed (22). Previous structural analyses on S protein revealed the receptor binding domains (RBDs) of D614G variant changed and its ability to bind to the receptor was improved (23,24). In another research, have been shown that D614G variant exhibits more competitive fitness and efficient infection in primary human epithelial lung cells. As well as transmission of this variant is significantly faster and then the wild-type virus (25).

The phylogenetic analysis demonstrated that most of circulating viruses in Iran belong to the lineage B.4 that is compatible with the other reports (8,26). B.1 is the most prevalent lineages in the world, although some reports from other countries show that other lineages are more frequent circulating variants (27). Furthermore, just as all other reports, we found that all the D614G mutations are related to the sequences belong to B.1 lineages (28).

In summary, in the current report we studied genomics variations and phylogeny of the SARS-CoV-2 sequences in Iran. Our results showed that most of the variations are related to the ORF1ab that are related to the virus replication and transcription. Also, we screened D614G mutation in spike protein and showed the variants with this mutation were introduced to Iran in May 2020 which can be the reason of increasing number of positive tests and deaths in this period of the time. Furthermore, we demonstrated that lineage B.4 is the most prevalent variant in Iran while B.1 lineage and the subsets including D614G mutation was detected in specimens sampled in May and after. As our data includes the specimens from March to September, there is an emergent need to analyze the viral lineage population with more samples and also beyond the September to screen the frequency of the B.1 lineage in comparison to the B.4. These data with similar future studies provide an opportunity to track and predict the transmission behavior patterns to apply appropriate strategy to control of the pandemic in Iran.

